# Operational and biochemical aspects of co-digestion (co-AD) from sugarcane vinasse, filter cake and deacetylation liquor

**DOI:** 10.1101/2021.02.24.432031

**Authors:** Maria Paula. C. Volpi, Antonio Djalma N. Ferraz, Telma T. Franco, Bruna S. Moraes

**Affiliations:** Interdisciplinary Center of Energy Planning, University of Campinas (NIPE/UNICAMP), R. Cora Coralina, 330 - Cidade Universitária, Campinas - SP, 13083-896, Brazil; School of Food Engineering, University of Campinas, Bertrand Russel, R. Josué de Castro, Cidade Universitária, Campinas - SP, 13083-000, Brazil; Centre for Environmental Policy, Imperial College London, Exhibition Road, London SW7 1NA, UK.; Chemical Engineering School, University of Campinas (FEQ/UNICAMP), Av.Albert Einstein 500, Campinas - SP, 13083-852, Brazil

**Keywords:** Co-digestion, 1G2G Sugarcane biorefinery, Methane production, continuous reactor operation

## Abstract

This work performed co-AD from the vinasse and filter cake (from 1G ethanol production) and deacetylation liquor (from the pre-treatment of sugarcane straw for 2G ethanol production) in a semi-Continuous Stirred Tank Reactor (s-CSTR) aiming to provide optimum operational parameters for continuous CH_4_ production. Using filter cake as co-substrate may allow the reactor to operate throughout the year, as it is available in the sugarcane off-season, unlike vinasse. A comparison was made from the microbial community of the seed sludge and the reactor sludge when CH_4_ production stabilized. Lactate, butyrate and propionate fermentation routes were denoted at the start-up of the s-CSTR, characterizing the acidogenic phase: the Oxidation-Reduction Potential (ORP) values ranged from -800 to -100 mV. Once the methanogenesis was initiated, alkalizing addition was no longer needed as its demand by the microrganisms was supplied by the alkali-characteriscs of the deacetylation liquor. The gradual increase of the applied Organic Load Rates (OLR) allowed stabilization of the methanogenesis from 3.20 gVS L^-1^ day^-1^: the highest CH_4_ yield (230 NmLCH_4_ gSV^-1^) and average organic matter removal efficiency (83% ± 13) was achieved at ORL of 4.16 gVS L^-1^day^-1^. The microbial community changed along the reactor operation, presenting different metabolic routes mainly due to the used lignocellulosic substrates. Bacteria from the syntrophic acetate oxidation (SAO) process coupled to hydrogenotrophic methanogenesis were predominant (∼ 2% Methanoculleus) during the CH_4_ production stability. The overall results are useful as preliminary drivers in terms of visualizing the co-AD process in a sugarcane biorefinery integrated to scale.

**Keypoitns:** Integration of 1G2G sugarcane ethanol biorefinery from co-digestion of its residues Biogas production from vinasse, filter cake and deacetylation liquor in a semi-CSTR Lignicellulosic substrates affected the biochemical routes and microbial community Biomol confirmed the stablismenht of thermophilic community from mesophilic sludge

## 1. INTRODUCTION

Anaerobic digestion (AD) of residues from sugarcane ethanol production has shown to be a promising strategy for waste management towards bioenergy enhancement (Janke et al. 2015; Moraes et al. 2015a; Janke et al. 2016). The most voluminous waste from the ethanol distillation columns, i.e, vinasse, has already shown potential for methane (CH_4_) production, even on a demonstration scale, reaching up to 310 NmL CH_4_ per gram of removed Chemical Oxygen Demand (COD) in a thermophilic Upflow Anaerobic Sludge Blanket (UASB) reactor from a Brazilian sugarcane mill in operation since the 1980s (Souza et al. 1992). The filter cake from the sugarcane juice filtration has also been considered a potential source for CH_4_ production as co-substrate in the AD process, with only one Brazilian company currently annoucing a co-AD technology, although scientific and widespread information is no longer provided (Zaparolli 2019). It is known that such residue is rich in trace elements with a favorable balance of macronutrients that can contribute to co-AD besides having a suitable average C:N ratio (24:1) for the AD (Janke et al. 2015). In this system, it is assumed that the AD could operate throughout the season without interruptions caused by the unavailability of vinasse in the off-season, using another liquid stream such as 2G vinasse or some other residue from the 2G ethanol pre-treatment steps.

The co-AD is an alternative process for AD of isolated substrates which may optimize the CH_4_ yield. It allows the use of residues with low biodegradability and/or inhibitory substances content by providing its dilution, apart from balancing micro and macronutrients and supplying synergistic effects between microorganisms (Hagos et al. 2017). It seems to fit in the management of residues from 2^nd^ generation (2G) ethanol production, which generates lignocellulosic waste usually recognized as complex substrates for AD. Within the co-AD concept, the integration of 1^st^ and 2^nd^ generation (1G2G) sugarcane biorefineries can be reinforced by blending their residues and maximizing their sustainable use to bioenergy generation.

Numerous types of pre-treatments of lignocellulosic biomass have been developing for the release of sugars (e.g., hexoses and pentoses) for the production of 2G ethanol (Moraes et al. 2015a). Alkaline pretreatments are common for the delignification of biomass, having additional effects on the silica removal (ash insoluble component) or the partial removal of hemicelluloses (including acetyl and uronic acid groups) and the swelling of cellulose, resulting in a substantial increase in the fiber surface (Carvalho et al. 2016).The residues generated are potential sources for AD (Rabelo et al. 2011), although little has been studied about the anaerobic co-AD, especially for the recent and innovative pre-treatment of biomass and hydrolysis, e.g., deacetylation process, pre-treatment with ionic liquids, hydrolysis using genetically modified yeast, among others (Nakasu et al. 2020). The complexity of such substrates for AD may be one of the factors driving the integration of the 1G2G ethanol process by co-AD of its residues, e.g., with 1G vinasse that is already recognized as a substrate for biogas production (Ferraz Júnior et al., 2016).

Previous studies by our research group were carried out concerning Biochemical Methane Potential (BMP) tests of waste from alkaline pretreatment of sugarcane straw (straw deacetylation) in co-AD with other residues from the sugarcane 1G ethanol mill. The results confirmed beneficial effects from the synergisms of the co-substrates (Volpi et al. u.d.). However, to the best of our knowledge, the behavior of the aforementioned waste from 2G ethanol production in semi-continuous bench-scale reactors has not yet been studied, aiming to provide preliminary basis for gradual scaling up of the process. Reactor operations should provide process parameters for continuous waste treatment and CH_4_ production, enabling us to forecast the maximization of residue use within their specific availabilities in the 1G2G sugarcane biorefineries.

AD stability and efficiency depend on the synergistic activity of the microorganisms that belong to the anaerbic consortium, which performs hydrolysis, fermentation, acetogenesis and methanogenesis activities (Li et al. 2009b). Relating the microorganism to its metabolic pathway is often a challenge. Identifying the microorganism in the process is already a big step, suggesting its metabolic potential, but it may not be enough to attribute the function of these microorganisms: a single microorganism may have different functions at different stages of the metabolic pathways (Cabezas et al. 2015). Furthermore, little has been found in the literature regarding the metabolic routes of microorganisms in AD from residues from the sugarcane industry with residues from 2G ethanol production in co-AD system.

Given this context, the objective of the present work was to perform the anaerobic co-AD of residues from 2G ethanol production (i.e., lignocellulosic liquor from sugarcane straw deacetylation pre-treatment) and 1G ethanol production (i.e., vinasse and filter cake) in a stirred bench-scale reactor with semi-continuous feeding. Monitoring the operation aimed to reach the upper limit of the organic load applied to the reactor for maximizing stable CH_4_ production, providing operational parameters for scale up of the co-AD process. Fundamental aspects of AD during operation were also investigated by the relation of reactor performance to monitoring analysis results. Microbial characterization was performed during stabilized CH_4_ production to relate the microorganisms to potential metabolic routes, as well as to assess the modifications in the microbial community from the seed sludge.

## 2. MATERIALS AND METHODS

### 2.1 Residues and Inoculum

The substrates were vinasse and filter cake from Iracema sugarcane mill (São Martinho group, Iracemápolis, São Paulo state, Brazil) and the liquor from the straw pretreatment process, performed at the National Biorenovables Laboratory (LNBR) from the Brazilian Center for Research in Energy and Materials (CNPEM). Deacetylation pre-treatment was applied to sugarcane straw on a bench-scale as described in Brenelli et al. (2020). The anaerobic consortium of the mesophilic reactor (BIOPAC®ICX - Paques) from the aforementioned Iracema mill was used as inoculum.

### 2.2 Semi-continuous reactor: description and operation

The semi-Continuous Stirred Tank Reactor (s-CSTR) consisted of a 5L-Duran flask with 4L-working volume, kept under agitation at 150 rpm by using an orbital shaking table Marconi MA 140. The operating temperature was 55°C, maintained by recirculating hot water through a serpentine. Thermophilic conditions were chosen because vinasse leaves the distillation columns at 90°C and thus would have lower (or none) energy expenditure to cool it to mesophilic conditions. Inoculum adaptation was performed by gradually increasing the temperature every 5 degrees per day until it reached 55°C, which was kept for a week before the beginning of the tests (Boušková et al. 2005). The pH adjustment to neutrality was performed by adding NaOH (1M) solution when necessary. The reactor was fed once a day with the blend of co-substrates (in terms of volatile solids, VS): 70% of vinasse, 20% of filter cake and 10% of deacetylation liquor, totaling 57.55 gVS L^-1^. These proportions were based on the residue’s availability at the sugarcane mill, where the most abundant is vinasse (25 L vinasse per liter of ethanol total (1G+2G)) and the least would be the deacetylation liquor (7 L per liter of ethanol total (1G+2G)). The reactor was fully filled with inoculum during the start-up, in which aliquots of effluent were discharged and new feed was added in a fed-batch mode of 24h throughout the operation. The Organic Loading Rate (OLR) applied to the reactor was increased over time to maximize the volume of treated waste with concomitant reduction of Hydraulic Retention Time (HRT). Table 1 presents the values of operational parameters applied to the s-CSTR according to the respective operation phases. Biogas volume and CH_4_ content were regularly monitored, as well as organic acids (OA), carbohydrates, alcohols, alkalinity and organic matter (in terms of VS) content in the digestate. Oxidation reduction potential (ORP) and pH were monitored both in the feed and digestate.

**Table 1.** Phases of reactor operation and the respective applied ORLs, feeding rate flows and HRT.

| Phase in Graph | OLR (gVS L<sup>-1</sup> day<sup>-1</sup>) | Feeding rate (L day<sup>-1</sup>) | HRT (days) |
| --- | --- | --- | --- |
| I | 1.50 | 0.100 | 40 |
| II | 1.80 | 0.125 | 32 |
| III | 2.30 | 0.160 | 25 |
| IV | 2.75 | 0.190 | 21 |
| V | 3.20 | 0.222 | 18 |
| VI | 4.16 | 0.285 | 14 |
| VII | 4.80 | 0.333 | 12 |
| VIII | 5.23 | 0.363 | 11 |

### 2.3 Analytical Methods

#### 2.3.1 Characterization of substrates

All the analyses followed the Standard Methods for the Examination of Water and Wastewater (APHA, 2012). The substrates were characterized in terms of chemical oxygen demand (COD) (method 5220B), series of solids (method 2540), pH (pHmeter PG 1800), OA, alcohol, carbohydrates, carbon, nitrogen and phosphorus (method 4500P). The COD was performed for the characterization of liquid substrates (vinasse and deacetylation liquor), using the digestion method and reading in spectrophotometer RAC DR 6000. Analyses of carbon, nitrogen and phosphorus were made using the TOC equipment Shimadzu-TOC-L-CNP. For the analysis of OA, carbohydrates and alcohols, the samples were centrifuged for 10 minutes at 10,000 rpm, filtered in porous membrane (0.2m m) and subjected to High Performance Liquid Chromatography (HPLC, Shimadzu®). The HPLC consisted of a pump equipped apparatus (LC- 10ADVP), automatic sampler (SIL-20A HT), CTO-20A column at 43 °C, (SDP-M10 AVP) and Aminex HPX-87H column (300 mm, 7.8 mm, BioRad). The mobile phase was H2SO4 (0.01 N) at 0.5 ml min^-1^. The series of solids included total solids (TS), volatile solids (VS) and fixed solids (FS), for all the substrates.

The elemental composition was performed for the characterization of filter cake in the Elementary Carbon, Nitrogen, Hydrogen and Sulfur Analyzer equipment (Brand: Elementar; Model: Vario MACRO Cube - Hanau, Germany).

#### 2.3.2 Monitoring of semi-Continuous Stirred Tank Reactor (s-CSTR)

Daily biogas production was measured using a Ritter gas meter, Germany. The CH_4_ content was determined by gas chromatography (Construmaq São Carlos) five times a week. The carrier gas was hydrogen (30 cm s^-1^) and the injection volume was 3 mL. The CG Column was made of a 3-meter-long stainless steel, 1/8” in diameter and packaged with Molecular Tamper 5A for separation of O_2_ and N_2_ and CH_4_ in the thermal conductivity detector (TCD). It had a specific injector for CH_4_ with a temperature of 350 °C, an external stainless-steel wall and an internal refractory ceramic wall. Detection (resolution) limits are 0.1 ppm for CH_4_.

VS analyses were also carried out during the reactor operation. The determination was performed in the feeding and digestate to account for the organic matter (in terms of VS) removed during co-AD. The pH and the ORP of digestate were measured, immediately after sampling (before feeding) using a specific electrode for Digimed ORP. Alkalinity was performed using the titration method (APHA, 2012). OA, carbohydrates and alcohol analysis was performed for digestate three times a week.

### 2.4 Calculations

Principal Component Analysis (PCA) was performed using STATISCA 10 through the correlation between the metabolites obtained (organic acids and alcohols) and the methane production, pH, partial and intermediate alkalinity, removal of organic matter and ORP variables.

Gibbs free energies (ΔG°) of the conversion of propionate to acetate were calculated at 55°C in pH 7.0. The values were computed in accordance with (Alberty 1998; Dolfing 2015).

### 2.5 Biology Molecular Analysis

Microorganism identification analyses were carried out for the seed sludge samples (Sample 1) before they were added to the reactor, and when the sludge was already stabilized in the s-CSTR with a stable production of CH_4_ under the OLR of 4.80 gVSL^-1^day^-1^ (Sample 2). Genomic DNA was extracted in triplicate and the PowerSoil DNA Isolation Kit (Mobio) was used. For visual confirmation of the quality and integrity of the DNA extracted from the samples, a run on a 1% agarose gel stained with SYBR® Safe (Invitrogen) was performed. DNA quantification in the sample was performed with the Qubit® 3.0 equipment Fluorometer (Life Technologies) and the quality based on the 260/280 ratio, which was determined using the Nano Drop Lite equipment (Thermo Fisher). The large-scale sequencing of the V3-V4 region of the 16S ribosomal RNA gene from bacteria and archaea present in the samples was then determined, in triplicate, using the Illumina MiSeq platform with paired-end sequencing (2 x 250 bp).

For the sequence analysis, the quality of readings was evaluated using the FastQC tool v.0.11.5 (Andrews, 2010), with quality strings lower than 30 (Phred score) and less than 100 base pairs were filtered with Trimmomatic 0.39 (Bolger et al. 2014). Bioinformatics analyses were performed using the Quantitative Insights into Microbial Ecology (QIIME2, version 2019.7, https://docs.qiime2.org/2019.7) (Bolyen et al. 2019) and its plugins. The taxonomic classification of Operational Taxonomic Units (OTUs) was performed with the q2-feature-classifier plug-in (Bokulich et al. 2018) in the classify-consensus-vsearch program (Rognes et al. 2016), based on the SILVA Ribosomal RNA Gene version 132 database (Quast et al. 2013). The resulting Qiime output file containing the abundances of OTUs in the samples was analyzed using the Phyloseq package (McMurdie and Holmes 2013) from the R (Team, 2013) software for making graphs and tables.

The large-scale sequencing of amplicons from the ribosomal operon of the microbial community led to identifying the bacteria and archaea present in the samples in-depth to characterize the microbiota. The results of the genera found were expressed in percentage, reflecting the relative abundance of microorganisms in the samples. Raw sequences were deposited in BioSample NCBI under accession number PRJNA684620.

## 3. RESULTS AND DISCUSSION

### 3.1 Characterization of residues

Tables 2 and 3 show the characterization of the inoculum and the residues fed to the s-CSTR. Two different batches of vinasse and deacetylation liquor were used throughout the operation, called Batch 1 and Batch 2. Batch 1 was used in the first stages of the operation and Batch 2 of vinasse and deacetylation liquor was fed from Phases IV and VII, respectively. The differences in these substrate compositions directly affected the reactor supply, making it necessary to adjust the feeding volume to maintain the applied OLR throughout the operation. The C:N ratios of vinasse Batch 1 (28:1) and Batch 2 (40:1) were in the recommendable range for AD processes (20 – 40:1) (FNR, 2010), although the C:N ratio was slightly higher in the latter, mainly due to its N content being about 64% lower than in Batch 1. The COD value of vinasse Batch 1 was close to the values normally found in the literature (Moraes et al. 2015a), whereas Batch 2 had much lower COD values. Accordingly, the level of TS was also lower than the vinasse in Batch 1, although the VS content was about 5 times higher. This fact shows the complexity of vinasse composition, which is significantly affected by factors such as the ethanol production processes and the sugarcane plant characteristics and cultivation. The choice of the suitable parameter for organic matter contabilization and its maintenance for the reactor monitoring directly affects the successful operation and consequently CH_4_ production. The COD of vinasse Batch 1 takes into account non-organic materials, e.g., sulfide from yeasts after the fermentation cycle as a way to prevent flocculation. In this study, these differences resulted in different vinasse volumes from Batch 1 and Batch 2 to compose the feed keeping the same applied OLR. Based on VS contents, larger volumes of vinasse from Batch 1 were used, which would be the opposite if only the COD was considered as a parameter for organic matter content. It highlights the importance of regular analysis for vinasse characterization throughout the season, especially related to the organic material, so that the applied OLR remains stable, thus avoiding organic load shocks, which can lead the reactor to failure (Fuess et al. 2017b). The “poor” operational control of vinasse AD normally adopted by Brazilian sugarcane plants results in an inefficient operation of the reactor, which has reflected in “negative marketing” for this process to scale in the sector. As the residue compositions vary throughout the sugarcane season and ethanol processing, the strict operation and monitoring of AD reactor and substrates are essential for success. Uncertainties regarding the production of 2G ethanol and its residues make it even more difficult to insert AD, and bench scale tests were extremely important to expand and deepen the knowledge of the main factors that affect this biological process.

**Table 2.** General Characterization of Residues

| Characterization | Inoculum <sup>a</sup> | Vinasse Lot 1 <sup>a</sup> | Vinasse Lot 2 <sup>a</sup> | Liquor Lot 1 <sup>a</sup> | Liquor Lot 2 <sup>a</sup> | Filter Cake <sup>a</sup> |
| --- | --- | --- | --- | --- | --- | --- |
| COD (g .L<sup>-1</sup>) | 11.171 ± 0.901 | 28.660 ± 0.91 | 17.020 ± 0.45 | 32.920 ± 0.27 | 10.440 ± 0.01 | - |
| Volatile Solids (g.mL<sup>-1</sup>) | 0.015 ± 0.001 | 0.018 ± 0.00 | 0.099 ± 0.00 | 0.103 ± 0.00 | 0.008 ± 0.00 | 0.192 ± 0.01 |
| Fixed Solids (g.mL<sup>-1</sup>) | 0.010 ± 0.002 | 0.007 ± 0.01 | 0.005 ± 0.00 | 0.021 ± 0.00 | 0.005 ± 0.00 | 0.097 ± 0.00 |
| Total Solids (g.mL<sup>-1</sup>) | 0.025 ± 0.020 | 0.025 ± 0.00 | 0.015 ± 0.00 | 0.011 ± 0.00 | 0.014 ± 0.00 | 0.290 ± 0.02 |
| pH | 7.350 ± 0.210 | 4.030 ± 0.34 | 4.090 ± 0.48 | 12.450 ± 0.13 | 11.450 ± 0.21 | - |
| Total Carbon (mg. L<sup>-1</sup>) | 4892.00 ± 0.11 | 14110.94 ± 0.21 | 12658.00 ± 0.02 | 15919.00 ± 0.89 | 16652.00 ± 0.33 | - |
| Organic Total Carbon (mg. L<sup>-1</sup>) | 3689.00 ± 0.58 | 14099.27 ± 0.12 | 12643.00 ± 0.45 | 15113.00 ± 0.56 | 15971.00 ± 0.56 | - |
| Inorganic Carbon (mg. L<sup>-1</sup>) | 1203.00 ± 0.69 | 11.69 ± 0.35 | 15.39 ± 0.63 | 805.60 ± 0.43 | 681.10 ± 0.33 | - |
| Nitrogen (mg. L<sup>-1</sup>) | 596.30 ± 0.73 | 495.93 ± 0.95 | 302.50 ± 0.15 | 176.30 ± 0.96 | 121.20 ± 0.28 | - |
| Phosphor (mg. L<sup>-1</sup>) | 13.72 ± 0.00 | 34.01 ± 0.00 | 9.96 ± 0.00 | 7.74 ± 0.00 | 2.98 ± 0.00 | - |
| Soluble Lignin (g. L<sup>-1</sup>) | - | - | - | 5.50 | 10.99 | - |

<sup>a</sup>Mean of three replicates ± standard deviation; -: not carried out

**Table 3.** Composition of Acids, Carbohydrates and Alcohols of residues

| Compounds | Vinasse Lot 1 | Vinasse Lot 2 | Liquor Lot 1 | Liquor Lot 2 |
| --- | --- | --- | --- | --- |
| Citric (mg L<sup>-1</sup>) | 237.99 | 0.00 | 0.00 | 0.00 |
| Succinic (mg L<sup>-1</sup>) | 278.33 | 205.06 | 0.00 | 0.00 |
| Propionic (mg L<sup>-1</sup>) | 1695.30 | 0.00 | 0.00 | 822.74 |
| Formic (mg L<sup>-1</sup>) | 1019.35 | 586.98 | 0.00 | 22.01 |
| Acetic (mg L<sup>-1</sup>) | 657.20 | 0.00 | 3670.00 | 1063.37 |
| Isobutyric (mg L<sup>-1</sup>) | 1304.90 | 216.43 | 78.84 | 0.00 |
| Butyric (mg L<sup>-1</sup>) | 160.30 | 549.95 | 0.00 | 0.00 |
| Malic (mg L<sup>-1</sup>) | 259.18 | 160.80 | 0.00 | 34.11 |
| Valeric (mg L<sup>-1</sup>) | 39.01 | 0.00 | 0.00 | 0.00 |
| Caproic (mg L<sup>-1</sup>) | 791.05 | 0.00 | 0.00 | 0.00 |
| Isovaleric (mg L<sup>-1</sup>) | 0.00 | 0.00 | 0.00 | 1788.62 |
| Lactic (mg L<sup>-1</sup>) | 3891.10 | 341.60 | 0.00 | 0.00 |
| Glucose (mg L<sup>-1</sup>) | 570.74 | 738.30 | 85.20 | 726.50 |
| Fructose (mg L<sup>-1</sup>) | 647.23 | 400.79 | 0.00 | 856.04 |
| Ethanol (mg L<sup>-1</sup>) | 114.30 | 0.00 | 0.00 | 0.00 |
| Butanol (mg L<sup>-1</sup>) | 556.62 | 0.00 | 0.00 | 0.00 |

Reinforcing the variability of the agroindustrial waste composition, phosphorus (P) content of vinasse in Batch 2 was much lower than that detected in Batch 1. The values are within the wide range reported in the literature (4-250 mg L^-1^) (Moraes et al. 2015b). P can be accumulated in phosphorus-accumulating organisms (PAO), in the starvation period, and be transformed into volatile fatty acids in anaerobic conditions from Polyhydroxyalkanoates (PHA) materials, potentially degraded into varied fractions of individual VFA according to the PHA composition; however, when these concentrations are in excess they can form buffer solutions that precipitate important minerals from AD such as calcium, magnesium, aluminum and iron (Wang et al. 2016), which was not observed in this work and, thus, P concentration was not inhibitory to the process.

Similar to vinasse, two batches of deacetylation liquor were used, in which Batch 1 contained higher COD, VS, TS, P concentrations than the liquor of Batch 2. On the other hand, soluble lignin content was twice as high in Batch 2 than in Batch 1. This compound was already reported to affect the AD process by its slow degradation, causing a “late” CH_4_ production (Mulat and Horn 2018). Low HRT applied to reactors may not take full advantage of the deacetylation liquor’s CH_4_ potential. Thus, co-AD reactors (normally CSTRs), known for their high HRT may be suitable for fully making the most of this substrate. The deacetylation liquor presented a C:N ratio of 90: 1 for Batch 1 and 137: 1 for Batch 2, showing insufficient N content against the C content. This reinforces the role of co-digestion for the balance of nutrients and dilution of components in excess, e.g., the liquor co-digested with vinasse and filter cake, both with higher levels of N.

Table 3 shows the main OA concentrations for the different batches of vinasse and deacetylation liquor, reinforcing the variability of such residues throughout the process and the season. Batch 1 of vinasse presented a larger variety of OA in higher concentrations, especially propionic acid, which can negatively affect the AD process (in concentrations as high as 900 mg L^-1)^ (Wang et al. 2009) by inhibiting the terminal acceptor – the methanogenic arqueas – and resulting in the accumulation of hydrogen (H_2_) and potentially raising the free energy (Marchaim and Krause 1993). Propionic acid was also detected in Batch 2 of liquor, in non-inhibitory concentrations. High concentrations of lactic acid were detected in Batch 1 of vinasse, and were reported to generate possible inhibition of AD because it is a precursor of propionic acid in the hydrolysis-acidification process (Zhang et al. 2007). Formic acid was also detected in vinasse Batch 1, which can be easily degraded by sulfate-reducing bacteria (SRB) (Dinsdale et al. 2000) and contributing to the sulfidric acid (H_2_S) generation in biogas. In the presence of sulfate, SBR competes with methanogenic archeas by the organic matter, leading part of the anaerobic metabolic pathways to sulfate reduction and lowering CH_4_ formation. The acetic acid content in both batches of liquor (especially Batch 1) indicates considerable potential for this residue to produce CH_4_ as the acetate pathway is the main one for CH_4_ formation (Lata et al. 2002). The presence and concentration values of the different OAs may lead to the predominant metabolic routes of AD process, which can change due to the variability of substrates composition and, thus, impairing the stable microbial consortia adaptation in AD reactors. Such complexity and specificities of these residues highlight the difficulty to introduce the co-AD process in continuous operation into the integrated 1G2G sugarcane biorrefiries, despite their considerable CH_4_ production potential.

The presence of isobutyric acid in vinasse Batch 1 and isovaleric acid in deacetylated liquor Batch 2 drew attention. These isoforms of such compounds have a worse rate of degradation in AD compared to their normal forms (butyric acid and valeric acid); however, the decomposition rate of the iso form of butyric acid is still higher than that of valeric and capric acid (Wang et al. 1999). Depending on the microbial consortia establishment, isobutyric acid can be degraded to acetic acid in the AD, which improves the CH_4_ production, or it can undergo reciprocal isomerization and become butyric acid. Isovaleric acid, on the other hand, does not undergo this reciprocal isomerization in the AD process, encompassing different little elucidated metabolic routes from that of valeric acid (Wang et al. 1999).

### 3.2 Semi-Continuous Stirred Tank Reactor performance

#### 3.2.1 Biogas production and reactor efficiency

Figure 1 shows the removal efficiency of organic matter and CH_4_ content in biogas produced in the reactor for each applied OLR. In the initial OLR (Phase I), large variations in the reactor efficiency occurred up to 55 days, when the organic matter removal stabilized at 71.27% ± 4.87% with the establishment of some metabolic routes for CH_4_ production (55.91 ± 5.78 NmLCH_4_ gSV^-1^). This behavior is in accordance with the results of digestate analysis (sections 3.2.2, 3.2.3 and 3.2.4). At each sequential increase of the applied OLRs, an initial disturbance on organic matter degradation was detected, representing firstly the adaptation of acidogenesis, at first 40 days (large variations on reactor efficiency) followed by the establishment of methanogenesis (little variations in the reactor efficiency). From about 90 days, CH_4_ production as high as 90 NmLCH_4_ gSV^-1^ was detected up to the penultimate applied OLR (Phase VII), which corresponded to the maximum applied OLR that the reactor was able to withstand with stability on CH_4_ production (93.92 ± 17.62 NmLCH_4_ gSV^-1^ and 79.57 ± 4.54 % of organic matter removal), although this was not the maximum CH_4_ yield. This fact indicates that methanogenesis activity started to become self-regulated from the end of Phase II (change of applied OLR from 1.80 gVS L^-1^day ^-1^ to 2.30 gVS L^-1^day ^-1^), which is reinforced by the results of digestate analysis presented in Sections 3.2.2, 3.2.3 and 3.2.4.

**Figure 1.**
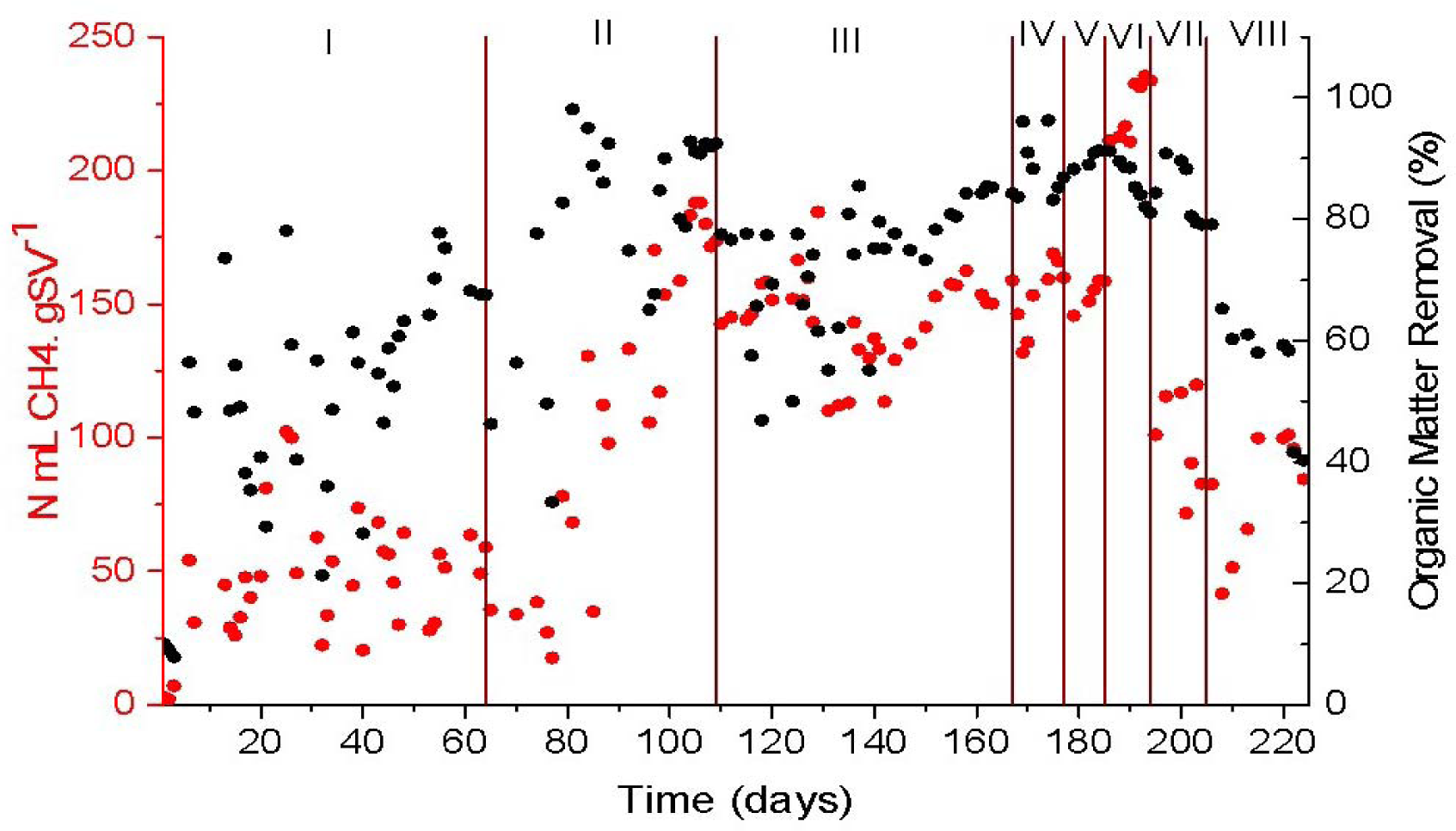
Methane production and organic matter removal along the reactor operation according to the applied OLRs (g VS L^-1^ day^-1^): 1.50 (Phase I); 1.80 (Phase II); 2.30 (Phase III); 2.75 (Phase IV); 3.20 (Phase V); 4.16 (Phase VI); 4.80 (Phase VII); 5.23 (Phase VIII)

The highest average reactor efficiency (83.08 ± 13.30 % organic matter removal and 233.20 ± 1.83 NmLCH_4_ gSV^-1^) and CH_4_ content (80.77% ± 0.28%) in biogas was achieved in Phase VI, corresponding to the specific biogas production of 324.85 ± 2.76 mL gSV^-1^. These values are close to those obtained by Janke et al. (2016) in which 320 ± 0.48 mL biogas g SV^-1^ was achieved at the maximum ORL of 3.0 g VS L^-1^day^-1^ in a s-CSTR treating sugarcane bagasse and filter cake (with tap water and cattle manure addition at mesophilic conditions, 38 °C). However, in their study, considerable OA accumulation in the digestate was observed (90% of OA, mainly propionic acid) and the average CH_4_ content in biogas remained about 50%.

Although a considerable decrease in CH_4_ content occurred at OLR 4.80 gVSL^-1^day^-1^ in the present work, CH_4_ production and reactor efficiency remained stable as already described. In a scale operation, the choice of the OLR to be applied and maintained in the reactor will depend on the objective of the operation: maximum volume of treated waste or maximum energy production in the form of CH_4_. The collapse of the studied s-CSTR occurred at the applied ORL of 5.23 gVS L^-1^day^-1^, when the reactor efficiency significantly dropped with the accumulation of OA (Figure 3A) and with the presence of carbohydrates in the digestate (Figure 3B)

#### 3.2.2 Degradation routes: pH and ORP indications

Figure 2 shows the monitoring of pH and ORP throughout the operation. In the first 40 days, the pH output was around 6 to 6.5 (Figure 2A), allowing the establishment of the acidogenesis process (Vongvichiankul et al., 2017), consistent with the starting behavior of the AD reactors, continuously adjusting the pH of the feed. After 90 days of operation, methanogenesis occurred and there was no need to adjust the pH, as the pH output was self-stabilized at around 7 until the end of the operation. ORP values followed the pH behavior, having larger variations (-800mV to -100mV) and less stable CH_4_ production (Section 3.2.1) during the beginning of the operation. Some studies reported considerable drops in the ORP values (-350mV to -550mV) during the period of the highest H_2_ production (Kataoka et al. 1997; Lin et al. 2008), corroborating the acidogenic step establishment at the beginning of the reactor operation. ORP variations during acidogenesis are related to the different metabolic pathways of acidogenic bacteria and the OLR applied to the reactor (Chen et al. 2015). The predominance of specific acidogenic routes may drive the methanogenic metabolic pathways, as well as the OA content of the fed substrates. After 90 days, the ORP variations decreased, although a considerable range still remained (-650mV to -450mV), which was further reduced (-300mV to -450mV) as CH_4_ production became more stable at the end of the operation. These values during stability are within the range reported as ideal conditions for acidogenesis and methanogenesis (Golkowska and Greger 2013).

**Figure 2.**
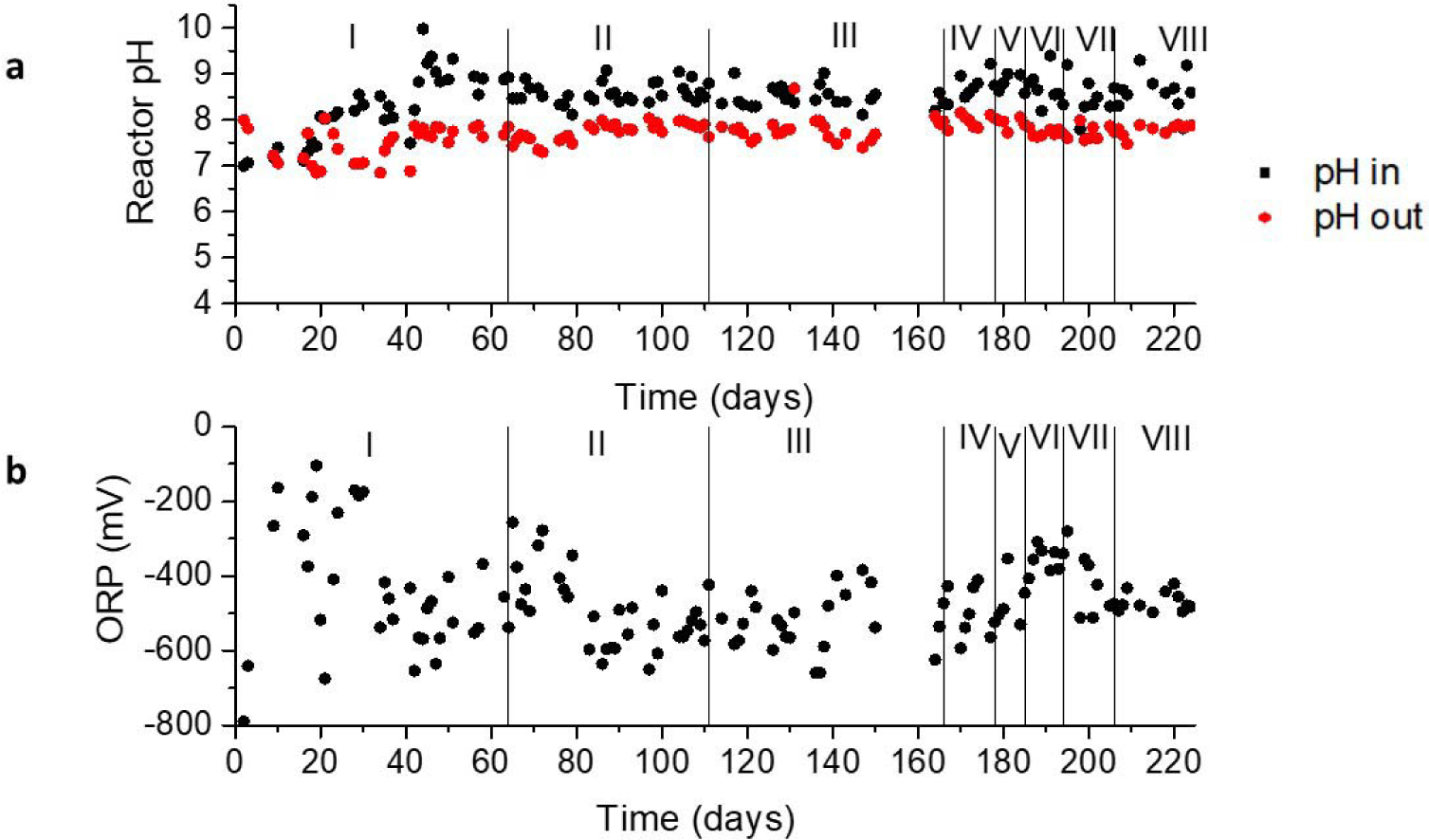
Monitoring of pH (A) and Oxidation Reduction Potential (ORP) (B) throughout the reactor operation according to the applied OLRs (g.VS.L^-1^.day^-1^): 1.50 (Phase I); 1.80 (Phase II); 2.30 (Phase III); 2.75 (Phase IV); 3.20 (Phase V); 4.16 (Phase VI); 4.80 (Phase VII); 5.23 (Phase VIII)

**Figure 3.**
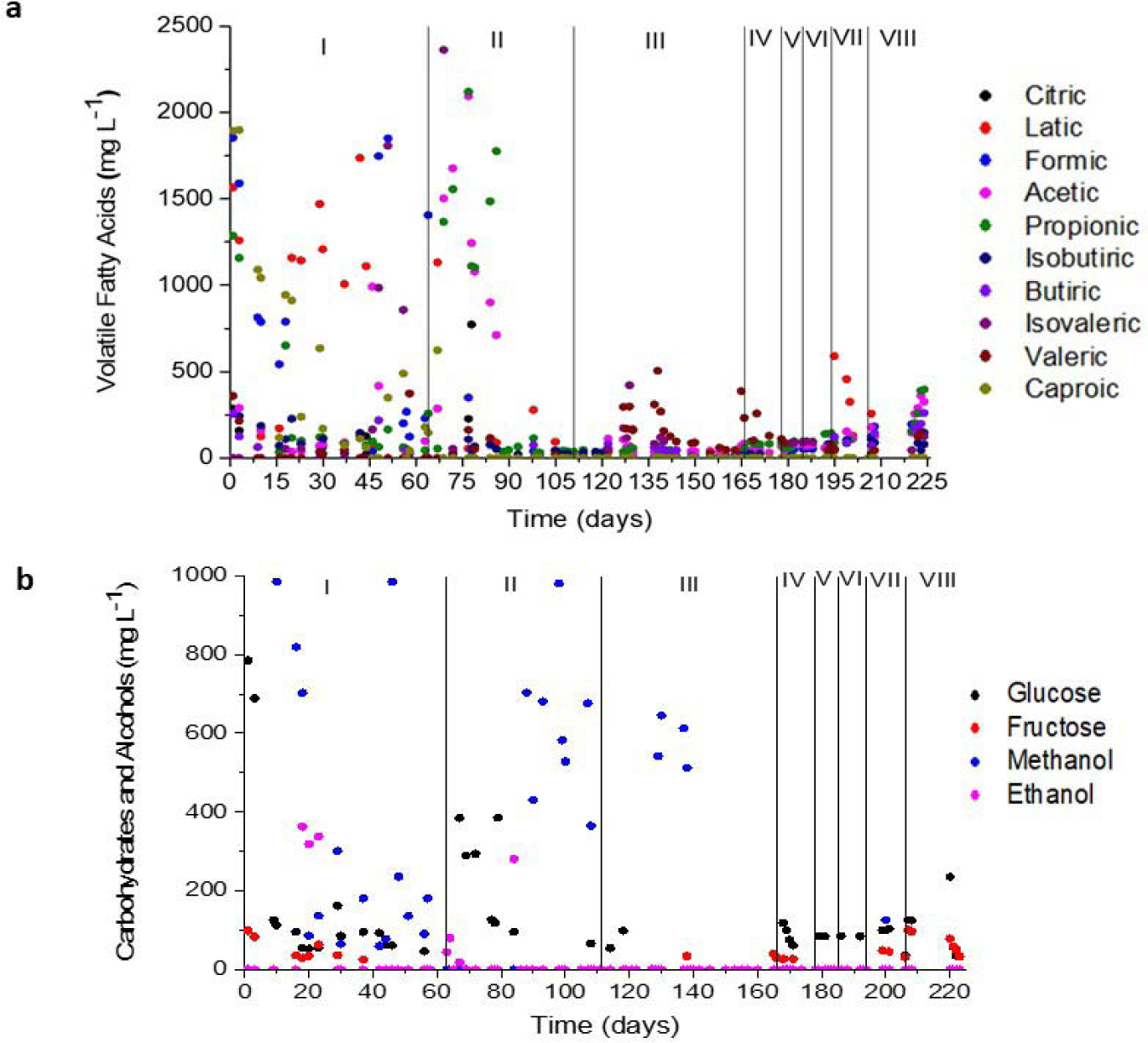
Values of (A) Organic Acids and (B) Carbohydrate and Alcohol concentrations in the digestate monitored along the reactor operation according to the applied OLRs (g VS L^-1^ day^-1^): 1.50 (Phase I); 1.80 (Phase II); 2.30 (Phase III); 2.75 (Phase IV); 3.20 (Phase V); 4.16 (Phase VI); 4.80 (Phase VII); 5.23 (Phase VIII)

In Phase II, when lactic acid was consumed, there were variations in ORP values, however the predominance was -600mV (Section 3.2.2). When the pH values of the outlet remained above 7 and ORP below -300mV, between 70 and 80 days (Figure 2A and Figure 2B - start of Phase II), the biogas production increased by 200%, and was even better after 90 days (392%), when methanogenesis was consolidated (Section 3.2.1). Vongvichiankul et al. (2017) also reported a considerable increase in the biogas production (from 1.88 to 22.90 L d^-1^) with the pH increase from 6.82 to 7.15 and the respective ORP increase from -359mV to -348mV.

#### 3.2.3 Degradation routes: OA, Carbohydrate and Alcohol indications

Figure 3 shows the results of the organic compounds monitored throughout the the reactor operation according to the applied OLRs. In the start-up OLR (Phase I), large amounts of OA were detected (Figure 3A), mostly from the residue composition itself (Table 3). According to the AD fundamental phases and the establishment of methanogenesis, these concentrations were decreased throughout the operation due to the conversion by acetogens into acetate, H_2_ and CO_2_, and then to CH_4_ by the methanogenic archeas to form biogas. The presence of the carbohydrates and their decrease in Phases I and II indicate the establishment of acidogenesis (Figure 3B). The consolidation of the methanogenic phase seemed to occur from about 90 days of operation, leading to a significant decrease in OA, alcohol and carbohydrates, although remaining methanol concentrations were detected. Methanol conversion in AD can occur by cultures of methanogens or SRBs. The methanogens convert methanol into methyl-coenzyme M and in the presence of hydrogen methyl-coenzyme, M is reduced to CH_4_ (Weijma and Stams 1999). When methanol is the sole substrate, however, part of the methanol has to be oxidized to CO_2_ to provide reducing equivalents for the reduction of methanol to CH4. This oxidation of the methyl-group likely proceeds via a reversed pathway which methanogens use to reduce CO2 to methane (Weijma and Stams 1999). In the presence of SRB, acetate is always necessary as a carbon source. The biochemical pathway of methanol oxidation by SRBs is not known. It is likely that methanol is oxidized to formaldehyde by means of a methanol dehydrogenase. Two other dehydrogenases oxidize formaldehyde to formate and then the formate is transformed into CO_2_ (Weijma and Stams 1999). As methanogens survive better in conditions of thermophilic temperature than SRBs, the methanogenic route from methanol should have been favored compared to sulfate reduction.

Lactic acid concentrations detected in Phase I, mostly from Batch 1 of vinasse composition, was probably converted into butyric acid (smaller proportions) or into propionic acid, both detected in Phase II, when lactate content significantly dropped (average ORP values close to -500mV and pH aroud 7). Chen et al. (2015) reported that butyrate-type fermentation can happen between -300 and -250mV, which is a lower range than obtained in the present study. The differences in substrate compositions and inocula may explain this fact, indicating that the microorganisms can adapt differently to the environment conditions. Sugarcane vinasse is a highly-acid substrate, and the butyrate-type fermentation is naturally favored by its composition (Fuess et al. 2020). On the other hand, Li et al. (2009) reported that ethanol-type fermentation is favored when there are high concentrations of acetic acid and ethanol. Chen et al. (2015) also detected the ethanol-type fermentation in the ORP value of -120 mV and the pH lower than 5. In the present study, during Phase II, acetic acid and small amounts of ethanol were also detected. It suggests the occurrence of butyric acid type fermentation (which may have been converted to acetate by SRBs) from the transition of Phase I to Phase II, with the ethanol type fermentation also taking place in the latter, even with the differences in the pH and ORP values reported in the few studies found in the literature, which reinforces the need for further research on this topic.

Although high content of malic and succinic acids were present in the fed substrates, no information regarding specific ORP values and their relationship with those acid-degradation metabolic pathways were found, but it is known they are propionate precursors in the AD process, as well as lactic acid (Scharer and Moo-young 1979). In Phase II, when metanogenesis started to be established, the lactate route may be shifted to propionic acid formation, as it was detected and sequentially consumed with the concomitant increase in CH_4_ prodution (Section 3.2.1). This situation may happen since the degradation of propionic acid is known to be the limiting factor in the CH_4_ production stage in thermophilic conditions (Djalma Nunes Ferraz Júnior et al. 2016). The ORP was close to -280 mV (begining of Phase II) and pH of 7.5, in agreement with the ORP data proposed by Wang et al. (2006) for this metabolic route, except for the pH (values reported close to 5.5). The main acid precursor of CH_4_, acetate, was detected up to 90 days of operation (ORP of -600mV), in parallel to the propionate appearance, when methanogenesis started to stabilize (Figure 2A). The slight delay on propionate consumption after acetate uptake occurred as the former is the last OA to stabilize due to its slow degradation rate (Wiegant et al. 1986)

Lactate formation was also observed in Phase VII, indicating that lactate producing-bacteria was established in the microbial consorcia of the s-CSTR. Fuess et al. (2018) also reported that this bacteria group played a role in the microbial dynamics of vinasse-fed acidogenic systems by providing an alternative carbon source for both H_2_-producing (butyric acid and H_2_ production) and non-H_2_-producing (propionic and acetic acids production) routes. With the organic overload of Phase VIII, the lactate to propionate route may have prevailed and the methanogens were not able to consume the latter acid, leading the reactor to collapse with a significant drop in the CH_4_ production and reomation organic matter efficiency (Section 3.2.1). It is worth mentioning that the increase of some carbohydrates (e.g., fructose and glucose) was also observed in Phase VIII, which suggests that the acidogenic step was also affected by the organic overload.

In Phase II, concentrations of propionic acid can be observed in the range of 1500 mg L^-1^ which is an inhibitory concentration for AD (Wang et al. 2009; Franke-Whittle et al. 2014). However, it did not inhibit the production of CH_4_ (Section 3.2.1), which can be explained by the fact that different systems have their own tolerance levels for OA due to the specific development and adaptation of the different microrganisms in the consortia to the diferent reactor conditions (Angelidaki et al. 1993). The decrease and stabilization of propionate concentrations along the reactor operation was a result of this self-regulation of anaerobic microbial consortium, avoiding the inhibition process by such acid accumulation. The stabilization of ORP values lower than the favourable one for propionic type fermentation (-278 mV) (Ren et al. 2007) also confirmed the minimization of this route.

A simplified thermodynamic analysis of the identified reaction 1 showed that there was no accumulation of propionic acid in the reactor, due to its conversion to acetic acid being favorable.

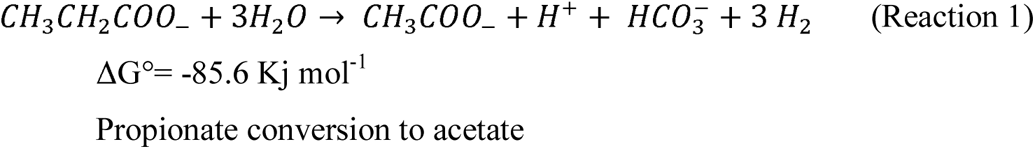

Zhao et al. (2018) showed that the higher temperature (thermophilic process) had positive effects on propionate acetogenesis, favoring its conversion to acetate. In addition, the literature shows that propionic acid degradation is better in systems with low H_2_ pressure and the concentration is kept low by H_2_ consuming methanogens (Wiegant et al. 1986). Hydrogenotrophic methanogens have been identified in the microbial consortium (Section 3.3) that may have contributed to the consumption of H_2_ and also favored the degradation of propionic acid when the methanogen stabilized (close to 90 days).

In Phase V, the low concentration of OA remained practically stable, with no major increases in OA up to Phase VI. In both phases, the reactor presented constant stabilization of CH_4_ production and removal of organic matter (Section 3.2.1). It confirms that the AD biochemical routes were self-regulated with the synergisms between acidogenesis and methanogeneses established.

#### 3.2.4 Degradation routes: Alkalinity indications

The alkalinity of the reactor (Figure 4) was in accordance with the behavior of pH, ORP and OA variables (Sections 3.2.2 and 3.2.3). Apart from the carbonate/bicarbonate system, the protonated forms of Volatile Fatty Acids (VFA) help to maintain the total alkalinity of anaerobic reactors, in which the intermediate alkalinity is caused by the ionized forms of VFA. The predominance of acidogenesis in the first 40 days of operation resulted in the low alkalinity values caused by the accumulation of VFA which is directly linked to the destruction of AD buffering capacity (Martín-González et al. 2013). In this period, the intermediate alkalinity/partial alkalinity (IA/PA) ratio remained between 1 and 2, much higher than the ideal value of 0.3, indicator of stability of AD process (Ripley et al. 1986). The gradual decrease in the IA/PA ratio close to 0.3 occurred over 90 days, coinciding with the consumption of the VFA’s (Figure 3A) and, thus, indicating the establishment of the self-controlled AD process. It is worth mentioning that alkalinizers were added only in the first days of Phase I, and were no longer needed in the following phases, indicating that the deacetylation liquor as co-substrate provided the necessary alkalinity for the AD system. Fuess et al. (2017a) showed that NaHCO3 is an alkalinizer used in AD processes, and is relatively expensive (USD 0.92 kg^− 1^) compared to the NaOH cost (USD 0.53 kg^-1^); therefore, the use of deacetylation liquor can further reduce costs. The suppression of alkalinizer use may represent an economic advantage for reactor operations, which could be decisive for the implementation of the AD technology.

**Figure 4.**
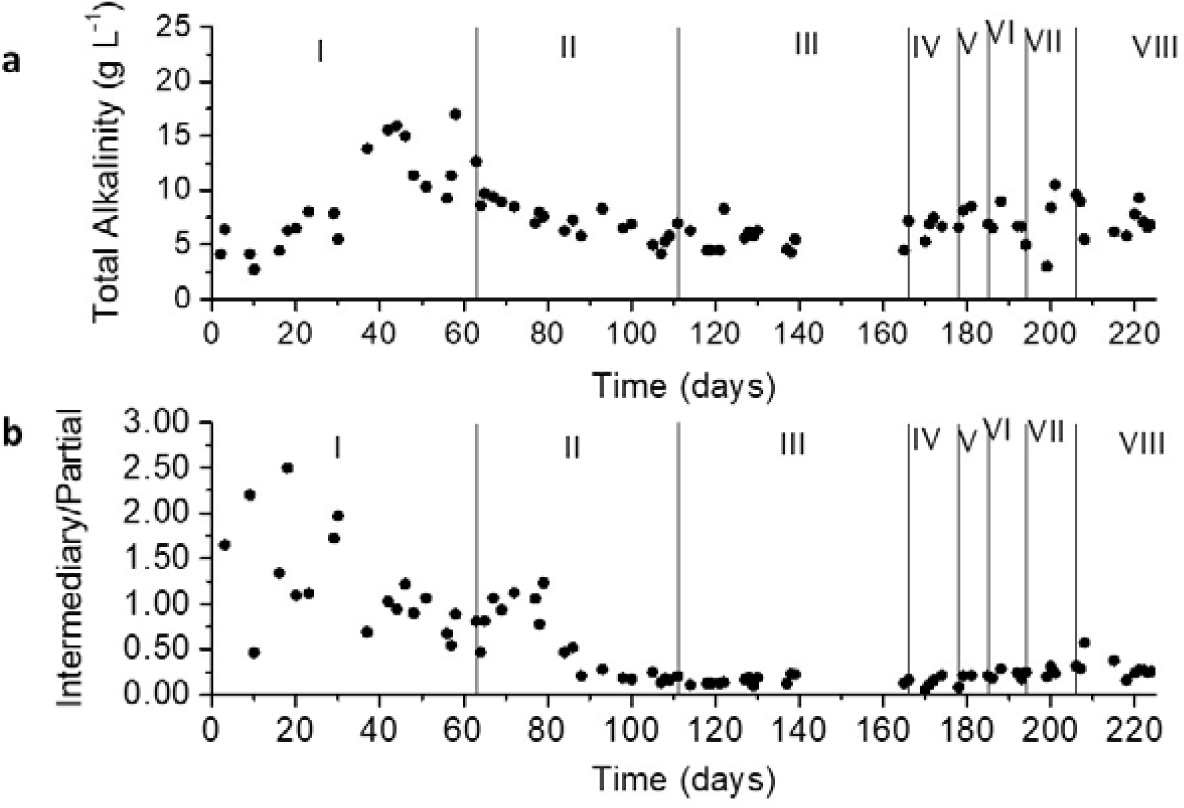
Monitoring of Reactor Alkalinity: (A) Total Alkalinity; (B) Alkalinity intermediary/Alkalinity Partial ratio according to the applied OLR’s (g VS L^-1^ day^-1^): 1.50 (Phase I); 1.80 (Phase II); 2.30 (Phase III); 2.75 (Phase IV); 3.20 (Phase V); 4.16 (Phase VI); 4.80 (Phase VII); 5.23 (Phase VIII)

#### 3.2.5 Relation of PCA and metabolic routes

Figure 5 shows the results of the PCA analysis, which was carried out to better understand the relationship of the metabolic routes with the variables of CH_4_ production, organic matter removal, alkalinity, ORP and pH.

**Figure 5.**
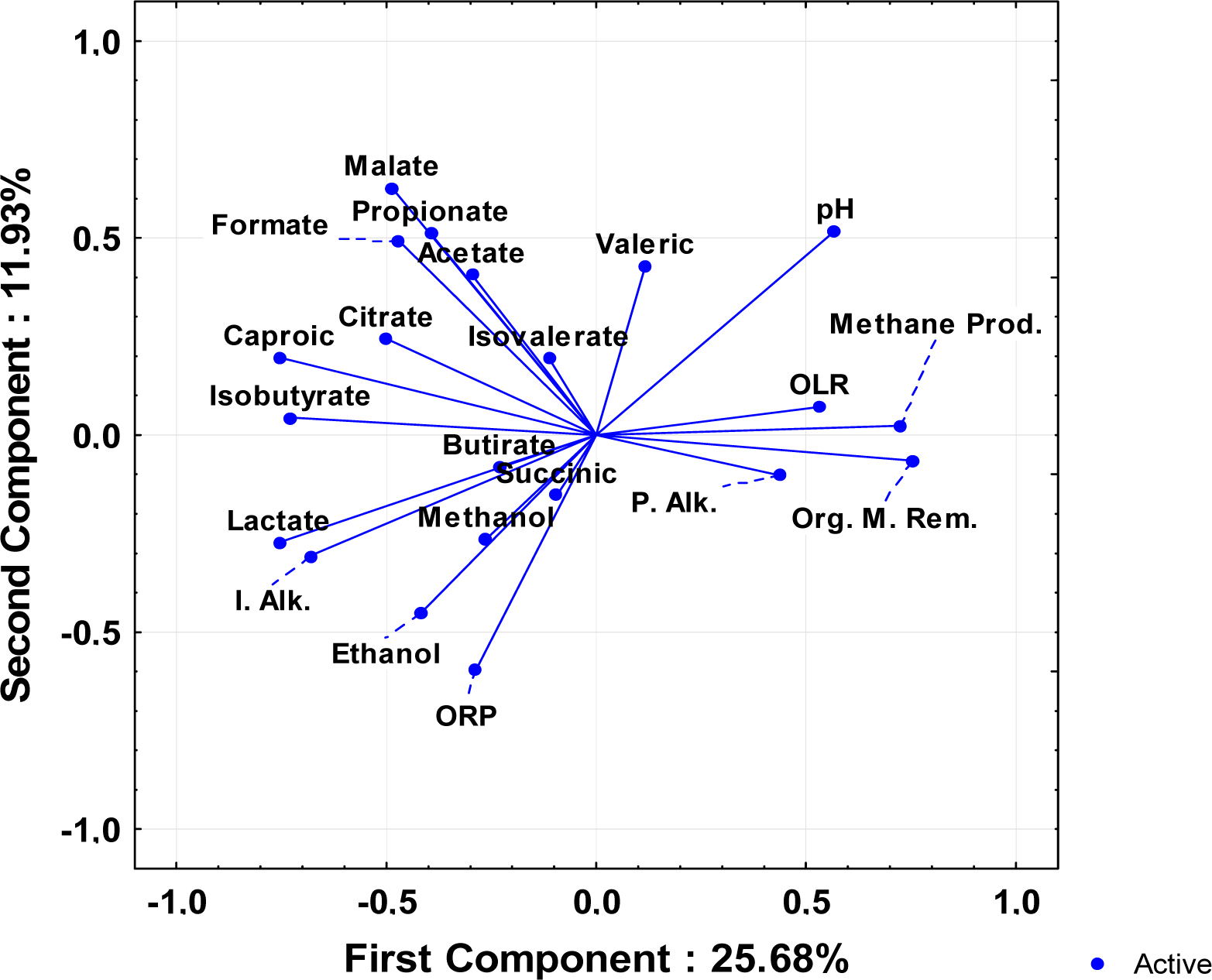
Principal components analysis of CH_4_-producing systems related with organic acids, pH, alkalinity and organic matter removal

Approximately 40% of the correlations can be explained by the PCA. The results showed that with the increase in OLR there was also an increase in CH_4_ production and consequently greater organic matter removal. These variables form a group and have an inverse relationship to the metabolites of lactic acid, methanol, ethanol, succinic acid, which makes sense, since the butyric-type fermentation can happen together with ethanol-type fermentation, in the acidogenic phase (Li et al., (2009)). In addition, lactic acid and butyric acid are precursors of propionic acid, also in the acidogenic phase (Krzysztof Ziemiński 2012).

The graph showed that the pH of the system, to favor CH_4_ production, needs to be closer to neutrality, since CH_4_ production is more related to partial alkalinity (5.75 <pH <8) than to intermediate alkalinity (4.3 <pH <5.75). In addition, intermediate alkalinity is indirectly related to pH, considering that low pH (4.3 <pH <5.75) is associated with reduced end-products such as lactate and solvents (ethanol, butanol and acetone) (Nunes Ferraz Júnior et al. 2020).

Through the analysis of PCA, it was also observed that as the concentration of OA (such as malic, acetic, propionic, formic acids) decreases, organic matter removal increases and consequently increases the production of CH_4_. This behavior is consistent with the results, since these acids are precursors of the phases of acidogenesis and acetoclastic methanogenesis (Vanwonterghem et al. 2015).

### 3.3 Microbial community characterization

Figure 6 shows the observed values of richness (number of species), and the calculated values from diversity (Shannon index) and wealth estimate (Chao1 estimator) of the Samples. The number of species (Figure 6A) and the richness (Figure 6B) of Sample 1 was greater than that of Sample 2. It is worth mentioning that the two Samples come from anaerobic reactors for CH_4_ production, varying the substrates and operating conditions, such as the temperature, in which Sample 1 comes from a mesophilic process and Sample 2 from a thermophilic process. This difference in temperature may have caused a selection of microorganisms, justifying these differences in the number of species and richness. In addition, operational and substrate differences may have led to CH_4_ production by different metabolic routes, selecting different microorganisms in the two Samples. Another reason that may explain this difference is that the microbial community in Sample 2, comes from a reactor stabilized in the CH_4_ operation, with the “selected” microorganisms.

**Figure 6.**
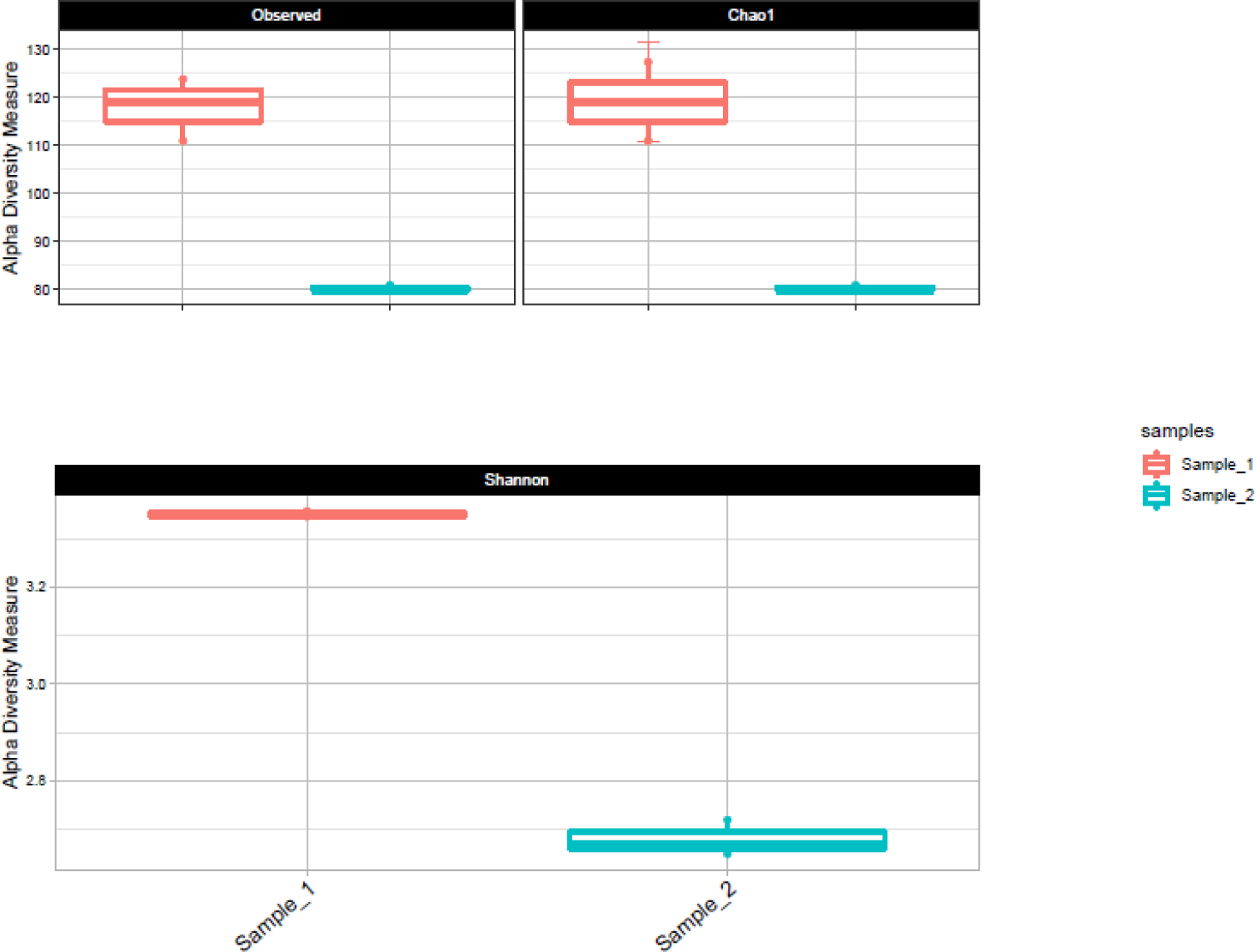
Observed values of (A) richness (number of species), (B) richness estimate (Chao1 estimator) and (C) calculated values of diversity (Shannon index) of Sample 1 (seed sludge) and Sample 2 (sludge from the s-CSTR stable opration, Phase VII)

Figure 6C shows the Shannon index, with values for both Samples below 4.0, which may indicate a greater specificity of the microorganisms. In general, when this index is greater than 5.0, it indicates a great microbial diversity in anaerobic digesters (Moraes et al. 2019). Even though both are below 5.0, Sample 2 still has a lower index, which indicates that the microbial community is even more specific, reinforcing the idea that these microorganisms are acting on different metabolic routes.

Figure 7A and Figure 7B show the relative abundances of microorganisms found in the Samples at the phylum and genus levels, respectively.

**Figure 7.**
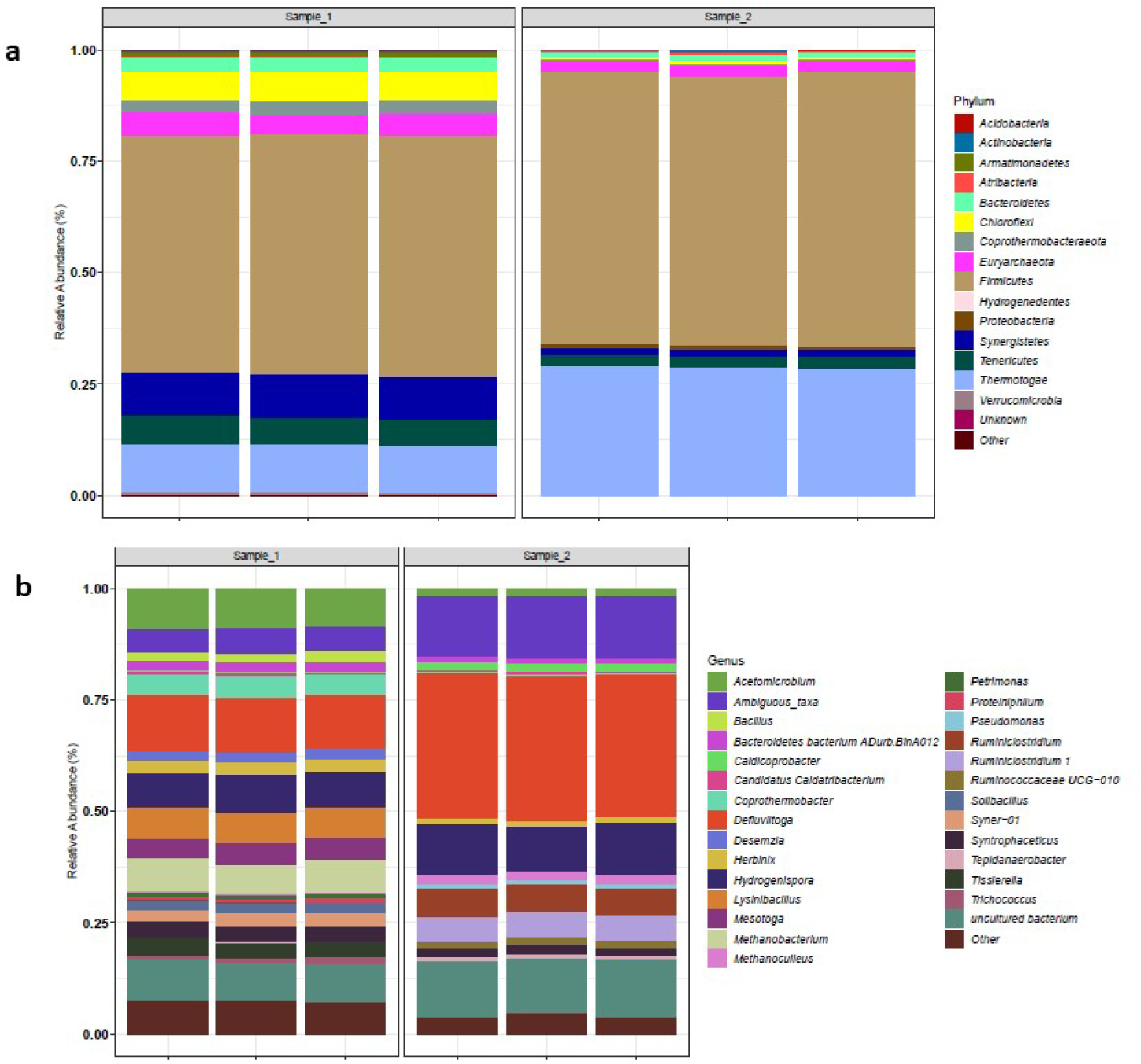
Relative abundance of microorganisms at the phylum level (A) and genus level (B) from the seed sludge- Sample 1 (1.1, 1.2 and 1.3) and from the s-CSTR sludge with stable CH_4_ production-Sample 2 (2.1, 2.2, 2.3)

Changes in the microbial community from one Sample to the other, both for phylum and genus, were observed. In Sample 1, there was a dominance of the microorganisms phyla: (∼3,5%) *Bacterioidetes*, (∼7,5%) *Chloroflexi*, (∼6,5%) *Euryacheota*, (∼50%) *Firmicutes*, (∼9%) *Synergistes*, (∼7%) *Tenericutes*, (∼11%) Thermotage. For Sample 2, the number of phyla was smaller, with greater abundance of: (∼53%) *Firmicutes*, (∼26%) *Thermotage* and (∼15%) *Euryacheota*. In both Samples, the relative abundance of the phylum Firmicutes was the highest, which is a common characteristic of the microbial community that makes up the anaerobic sludge (Chen et al. 2016; Wu et al. 2020). Bacteria from the phylum Firmicutes are the main ones that produce cellulosic enzymes within thermophilic AD. This phylum also contains acetogenic bacteria that degrade OA to produce acetic acid (Yu et al. 2018). The presence of the phylum *Firmicutes* may also be related to lignocellulosic residues as substrate (deacetylation liquor and filter cake in this work), as reported by Yu et al. (2018) using rice straw in thermophilic AD process. A decrease in the relative abundance of the phylum *Synergistes* and *Tenericutes* from Sample 1 to Sample 2 and an increase in the Phylum Thermotage in Sample 2 was also observed. The phylum *Thermotage* is the predominant one in thermophilic processes (Wang et al. 2018) and it has often been reported in thermophilic digesters treating organic wastes such as swine slurry, market biowaste, and food wastewater (Kim et al. 2018).

The Phylum *Proteobacteria* (∼0.2%) is related to the degradation of lignocellulose in the hydrolysis phase (Yu et al. 2018; Wu et al. 2020), and this phylum is observed only in Sample 2. The appearance of this phylum, even in small relative abundance (probably because the deacetylation liquor and filter cake were the co-substrate in minor proportions), indicates how the microbial population changes with the presence of different substrates for the CH_4_ production.

The detected groups of methanogenic microorganisms belong to the phylum Euryacheota (Wu et al. 2020), and are more abundant in Sample 1. However, Sample 2 presented different genera of methanogenic microorganisms comparatively to Sample 1: Methanoculleus (∼2%) and Methanobacterium (∼9,5%) in the respective Samples. Microorganisms from the aforementioned genus are hydrogenotrophic methanogen, using mainly H_2_ and CO_2_ for conversion to CH_4_ (Krzysztof Ziemiński 2012). The dominant methanogens in Sample 2 (Methanoculleus genus) have been reported to be predominant in the biogas production of mesophilic reactors with syntrophic oxidation of acetate (SAO) coupled with hydrogenotrophic methanogenesis (Schnürer et al. 1999). On the other hand, Hattori, (2008) reported that SAO coupled with hydrogenotrophic methanogenesis can also occur at elevated temperatures, and is confirmed by analyses in thermophilic digesters. Thus, our results indicate that CH_4_ was mainly produced from acetate via H_2_ and CO_2_ reduction in the s-CSTR, which implies the syntrophic oxidation of acetate coupled to hydrogenotrophic methanogenesis.

*Syntrophaceticus* and *Tepidanaerobacter* are known as SOA microrganisms which can also be coupled to hydrogenotrophic methanogenesis (Kim et al. 2018). Their presence in Sample 2 (∼2% and ∼0.5%, respectively) confirms the possibility of methanogenesis occurring from acetate via H_2_ and CO_2_ reduction, and these microrganisms are involved in acetate catabolism. The SOA coupled to the hydrogenotrophic methanogenesis metabolic route was observed in AD processes from residues with high protein and ammonium contents such as chicken manure, but also in reactors with *Jatropha* press cake as substrate (Ziganshin et al. 2013), that have lignocellulosic characteristcs such as the filter cake, i.e., a co-substrate used in the present work.

The predominance of the genus *Defluviitoga* (∼36%), belonging to the phylum *Thermothage*, was also observed in Sample 2. *Defluvitoga* genus is reported to be dominant in the degradation of organic materials in CSTRs or thermophilic bioelectrochemical reactors (Guo et al. 2014). Some members of the aforementioned genus can also metabolize sugars to generate H_2_ and OA in the hydrolysis phase of AD. This suggests that these microorganisms may also interact with hydrogenotrophic methanogenic microorganisms (Kim et al. 2018).

A relative abundance of the genus *Ruminoclostridium* (∼5%), belonging to *Firmicutes* phylum, was also observed in Sample 2, while it was not detected in Sample 1. Some species of the *Ruminoclostridium* genus are characterized by having acetate as their final product of sugar metabolism. They are known to metabolize materials with high concentrations of cellulose due to the high production of cellulolytic enzymes (Badalato et al. 2017). Studies by Peng et al. (2014) showed that bacteria of the *Ruminoclostridium* genus improved the efficiency of CH_4_ production using lignocellulosic residues, such as wheat straw as substrate. The presence of this group of microorganisms in Sample 2 indicates they were acting in the hydrolysis phase of cellulosic and lignocellulosic residues used in the co-digestion, thus explaining the reason for their absence in the seed sludge (Sample 1).

It is worth mentioning that the biomolecular analysis was performed only at Phase VII (OLR 4.80 gVS L^-1^day ^-1^), when the thermophilic methanogenesis was already established with stability, which is difficult to relate to the different transition metabolic routes mentioned in Section 3.2.3 about the fermentation of butyric, lactic and propionic acids. Detman et al. (2018) reported that lactic acid is oxidized to acetate via acetotrophic methanogenesis, and the main methanogenic is *Methanosarcin*. However, our results indicated the predominance of the SAO route, with hydrogenotrophic methanogenic organisms (Sample 2), implying the possibility that in Phases I and II of the reactor operation, there was a presence of acetotrophic microbial communities and SRB (such as *Desulfobulbus*) (Section 3.2.3), which have lactate utilization genes (Detman et al. 2018), but in Phase VII there has already been a change in the microbial community due to methanogenesis stability.

## 4. CONCLUSION

The co-AD in the s-CSTR proved to be a suitable alternative for energy recovery from the 2G ethanol production waste coupled to the residues from the 1G process. The reactor operation was effective for providing process parameters for continuous waste treatment and CH_4_ production, enabling us to forecast the maximization of residue use within their specific availabilities in the 1G2G sugarcane biorefineries. The upper limit of the OLR applied without collapsing the reactor was 4.80 gVS L^-1^day^-1^, maintaining the efficiency of the reactor and the stability of CH_4_ production, although being 59% lower than the maximum CH_4_ yield obtained at a lower OLR. These findings can guide practical operations in a biorefinery according to their current demand for energy production or maximizing waste treatment, changing the OLR applied to the reactor based on basic empirical science.

The composition of substrates played a role in establishing the predominant metabolic routes into the s-CSTR: lactate and butyrate degradation pathways seemed to mostly occur due to the high content of such acids in vinasse and deacetylation liquor. The liquor composition also contributed to keeping the reactor buffer capacity as its alkaline characteristics favored the lower addition of alkalinizers along the operation: it could result in cost reductions in an industrial scale as such co-substrate could partial or totally replace the alkalizers demand.

The change in the microbial community from the seed sludge and from the reactor sludge when CH_4_ production stabilized confirmed that the substrates composition and the operational conditions significantly affect the metabolic pathways for CH_4_ production. The biomolecular analysis results proved the feasibility of the establishment of thermophilic methanogenic community (∼26% *Thermotage*) from the mesophilic sludge. The specific substrates and reactor conditions directed the co-AD process, enabling the use of specific and already adherent microbial communities, or favoring the development of new related ones.

## ACKNOWLEDGEMENTS

This work was supported by FAPESP Project (2018/09893-1), FAPESP project (2016/16438-3) and FAPESP-BBSRC (2015/50612-8). The authors gratefully acknowledge the support of the Laboratory of Environment and Sanitation (LMAS) at the School of Agricultural Engineering (FEAGRI/UNICAMP), the National Laboratory of Biorenewables (LNBR/CNPEM) and the Interdisciplinary Center of Energy Planning (NIPE/UNICAMP).

## AUTHOR CONTRIBUTION STATEMENT

MPCV: Conceptualization, Investigation, Methodology, Data curation and Writing-original draft preparation. ADFNJ: Methodology, Resources, Data curation and Writing-original draft preparation. TTF: Project administration and Funding acquisition. BSM: Conceptualization, Formal analysis, Writing - Review & Editing, Supervision and Funding acquisition. All authors read and approved the manuscript.

